# A rapid and inexpensive universal PCR protocol for DNA (meta)barcoding using a one-tube, 2-step PCR

**DOI:** 10.1101/2025.05.16.654411

**Authors:** Olivier Collard, Mohammed M. Tawfeeq, Hugo Ducret, Jean-François Flot

## Abstract

DNA barcoding has become a widespread technique for species identification through the amplification of standardized gene regions. These include the cytochrome c oxy-dase subunit I (COI) for metazoans, ITS for fungi and plants or the 16S rRNA for bacteria and archaea. It has been shown to be more accurate than more traditional methods such as morphology, making it a now widespread method for species delimitation. The development of new sequencing methods such as Nanopore sequencing becoming more affordable and accurate has led to a growing use of those methods and a need for fast, standardized and cheap protocol. Although Sanger sequencing has become superseded by next-generation sequencing for most genomic applications, using second or third-generation sequencing for amplicons remains an expensive and time-consuming approach, especially for laboratories that analyse many different markers. Amplicon sequencing typically occurs in the context of either DNA barcoding (sequencing one or a few markers of interest for each specimen in a collection) or DNA metabarcoding (investigating the composition of a community by using universal primers targeting a whole group of organisms). Pooling the resulting PCR products for e.g. Nanopore sequencing requires the use of barcodes that are generally added to amplicons once they are amplified (using either ligation or a second round of amplification), and these supplementary steps are time-consuming, expensive and prone to contamination. To alleviate these issues, we present here a quick, single-tube two-step PCR with four primers: two internal primers with tails, and two external barcode primers that anneal on the tails during the second phase of the PCR. we show an improved 4 primers PCR protocol by using a single tube “drop in the lid” method. 8 DNA extract from *Asellus. sp* were obtained and amplified using both, 2-step PCR and the 1-step PCR. Results shows a clear enhancement of PCR product using the 2-step PCR for both ITS and CO1 marker compare to the 1-Step protocol.

**Method Summary:** To maximize the cost and effectiveness of barcoding approaches, we developed barcodes based on the Native Barcoding Kit 96 V14 of Oxford Nanopore Technologies (ONT). This step allows to tag samples within the PCR rather than through the use of ligase enzymes. To avoid primer interaction between the inner primer and the barcodes, the latter were added within the PCR cycles through the implementation of a hold step between the inner primer cycles and the barcoding cycles. For this purpose, barcodes are dried on the lid, avoiding the opening PCR tubes and preventing potential contamination.

## Introduction

Since its initial development in the 1990s, DNA barcoding has become a widespread method among scientists from different fields. Its use ranges from taxonomic identification (Frézal and Leblois, 2008; Hebert and Gregory, 2005) to phylogeography and community ecology (Kress et al., 2015). To date, most DNA barcoding techniques involve the use of costly devices (Illumina, Sanger) that can be difficult to access and require expensive investment or constant samples shipping. Next Generation Sequencing (NGS) tools such as those developed by Oxford Nanopore Technologies (ONT) can tackle these issues due to their affordable costs and user-friendly interfaces.. However, ONT sequencing often requires the constant use of dedicated kits involving expensive ligases, which can make the cost comparable to older methods.

By using 2 different annealing temperatures, with a lower annealing for the inner primers and an upper annealing for the barcodes, recently developed 4-primers PCR allow for simultaneous amplification and barcoding through one single PCR (Matsuo et al., 2021). These methods allow to amplify and barcode the targetted marker with reduced contamination and dimers production. However, the affinity between the inner primers and the DNA can significantly influence the effectiveness of the PCR. Therefore, with the introduction of barcodes during the PCR cycles, the interaction between the inner primers and the barcodes can be minimized. To prevent this contamination prone manipulation, barcodes can be dried on the PCR lid allowing their introduction at a different time period without the need of opening the PCR tubes (Macedo Rafael De Arruda et al., 2024).

Overall, the two-step PCR is composed of two separated PCR cycles (Figure 1). The separation is done by adding a 10°C hold step within the PCR cycles, putting the thermocycler on pause, allowing for tube manipulation. First, the DNA samples are briefly amplified with the inner primers for up to 5-20 cycles at a low annealing temperature. Second, barcodes previously dried on the PCR tube lid (following the protocol of Macedo Rafael De Arruda et al., 2024) are added into the PCR tube by shaking. The PCR is then resumed for 20-35 higher annealing barcoding temperature. Finally, For the sequencing, PCR product are cleaned using beads to remove primers dimers and PCR waste. Sequencing is then performed using Ligation sequencing amplicons V14 (SQK-LSK114) and a Flongle flow-cell.

**Figure 1.**
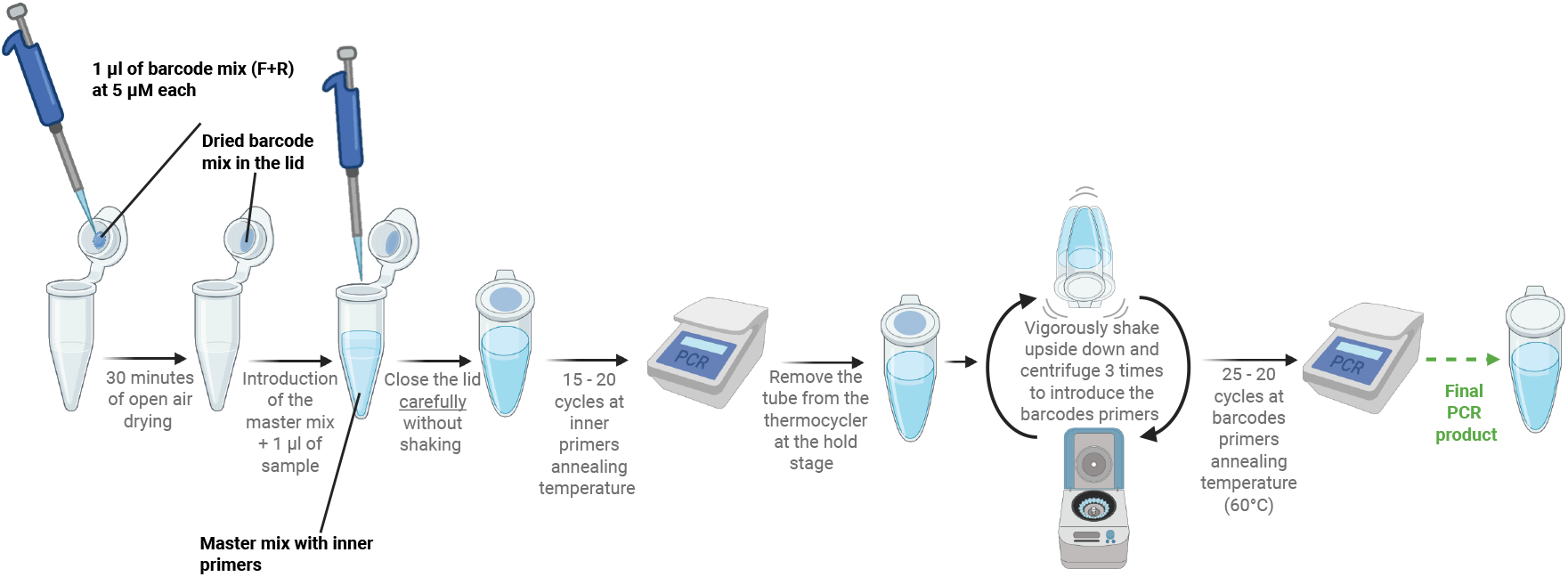
Adapted diagram of the “drop in the lid” method devlopped by Macedo Rafael De Arruda et al., 2024. The barcodes are represented in purple, and the master mix is shown in light blue. Created in BioRender. Flot, J. (2025) https://BioRender.com/x2r02dy

The aim of this work is to test this improved 4 primers PCR by comparing it with the traditional protocol. We compare the PCR yield obtained by simply performing the PCR with the 4 primers as described by Matsuo et al., 2021 (One-step PCR) with the enhanced “drop in the lid” method described by Macedo Rafael De Arruda et al., 2024 (Two-step PCR).

## Material & methods

In this study, 8 specimens of *Asellid* isopods were collected from wells in Avram Iancu (Romania). The samples were preserved in absolute ethanol and the tubes were kept at -20°C until Extraction. DNA extractions were then performed using Nucleospin Tissue extraction kit. For one-step PCR: the mix consisted of 6 *μ*l 2x Phire taq polymerase PCR MM (ThermoFisher Scientific, USA), 0.5 *μ*l of each primer (10*μ*M), 0.5 *μ*l DNA template and 4.5 *μ*L of nuclease-free water for a total volume of 11 *μ*L, 0.5 *μ*l of the forward and 0.5 of reverse barcodes were dried on the PCR tube cap then the tubes were shaken vigorously to mix the barcodes with the PCR mix prior placing them in the thermocycler (see figure 1). A thermocycler was then used with the following settings for the cytochrome oxidase 1 (CO1) marker: initial denaturation at 95°C for 300s, 17 cycles of 98°C for 8s, 53°C for 8s, 72°C for 20s. Following those cycles, 30 cycles of 98°C for 8s, 60°C for 8s, and 72°C for 20s, followed by a hold step at 10°C. The same settings were used for the internal transcribed spacer (ITS2), except that the settings of the initial denaturation step were changed to: 17 cycles of 98°C for 8s, 50°C for 8s, 72°C for 20s. For two-step PCR, we used the same PCR protocol except that we made a hold step before the start of 30 cycles, to mix the PCRs with their barcodes, centrifuge them, and place them back in the thermocycler. For both PCR protocols, the lid cover temperature was kept at 105° for all PCR runs. The inner primers used are described in table 1 and barcodes used are the one [provided by nanopore](https://nanoporetech.com/document/ligation-sequencing-gdna-native-barcoding-v14-sqk-nbd114-96) to which the linker was added.

**Table 1.**
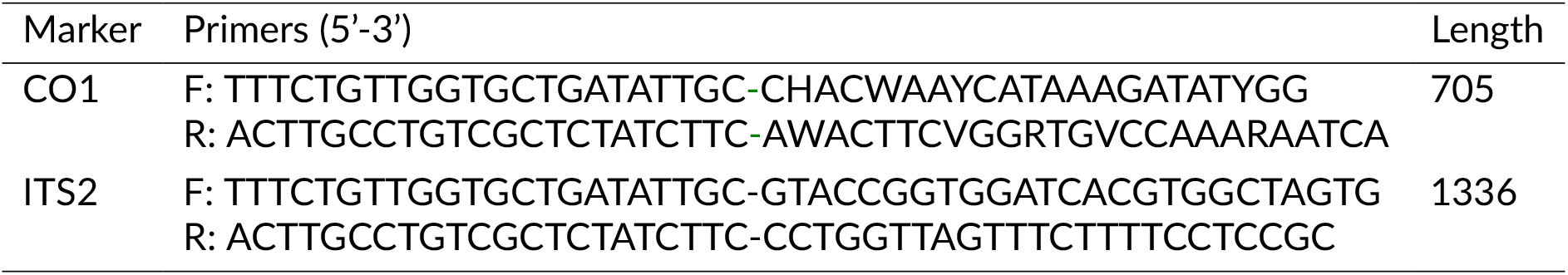
Nucleotide sequences of PCR primers. Nucleotides in yellow represent the tail, while nucleotides in green represent the primer sequence. The ITS primers were retrieved from Grutter et al., 2000, and the CO1 primers from Flot et al., 2010

A 1% agarose gel was prepared by mixing 1.2 g of agar-agar in 100 ml of TBE (45 mM Tris-borate, 1mM EDTA). The mixture was then melted using a microwave and cooled for 10 minutes. 5 *μ*l of Midori Green Advance was then added in the gel and poured in an electrophoresis tank before letting it solidify for 30 minutes. 4 *μ*l of samples were then added in each well followed by a migration of 20 minutes at 150V. The gel was then placed under UV for analysis. Following the PCR, products were polled in a single Ependorf tube with different ratio based on the thickness of the band observed on the gel. The pool was then cleaned using SPRIselect Bead-Based (Beckman) with a 0.65 ratio. PCR pool was then sequenced using Ligation sequencing amplicons V14 (SQK-LSK114) and a flongle flow-cell. Sequences were bascalled and demultiplexed using Guppy and consensus were made using the pipeling Amplicon_sorter (https://github.com/avierstr/amplicon_sorter).

## Results

The amplification of the ITS2 and CO1 markers using the two different methods (one-step and two-step PCR protocols) has shown very distinct results despite having the same Master Mix preparation protocol and PCR cycle.

For the ITS2 marker (Figure 2), the one-step PCR protocol displays an extremely faint band at the target sequence length (1336 bp) for sample 8 while showing no amplification for the other specimens. Alternatively, the two-step PCR protocol shows a significant increase in amplicon amplification compared to the one-step protocol. Both methods produce a substantial amount of unavoidable primer dimers due to the complementarity of the tail sequence present in both the inner primers and the barcode (see Table 1) Nevertheless, these small fragments can be easily removed using a selective cleaning protocol, such as the [SPRIselect Beads-based protocol](https://www.beckman.com/reagents/genomic/cleanup-and-size-selection/size-selection/spriselect-protocol)

**Figure 2.**
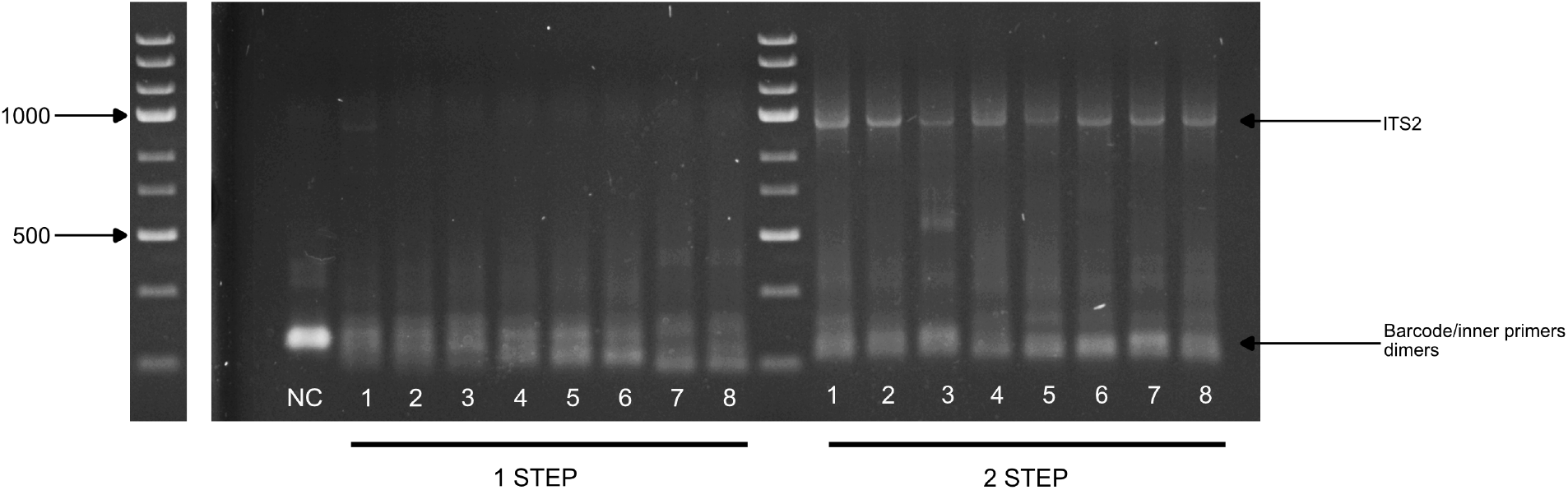
PCR comparison between the two-step (bottom) and one-step (top) protocols for the ITS2 marker. Samples are numbered from 1 to 8, each representing a different specimen of Asellus sp. Lane M: molecular size markers. Molecular size (in base pairs) is indicated on the left.

The CO1 marker (Figure 3) shows a similar result to the ITS2 marker, except for the absence of any amplification with the one-step PCR. The amplification for all specimens is significantly increased with the two-step PCR, along with a decrease in primer dimer production.

**Figure 3.**
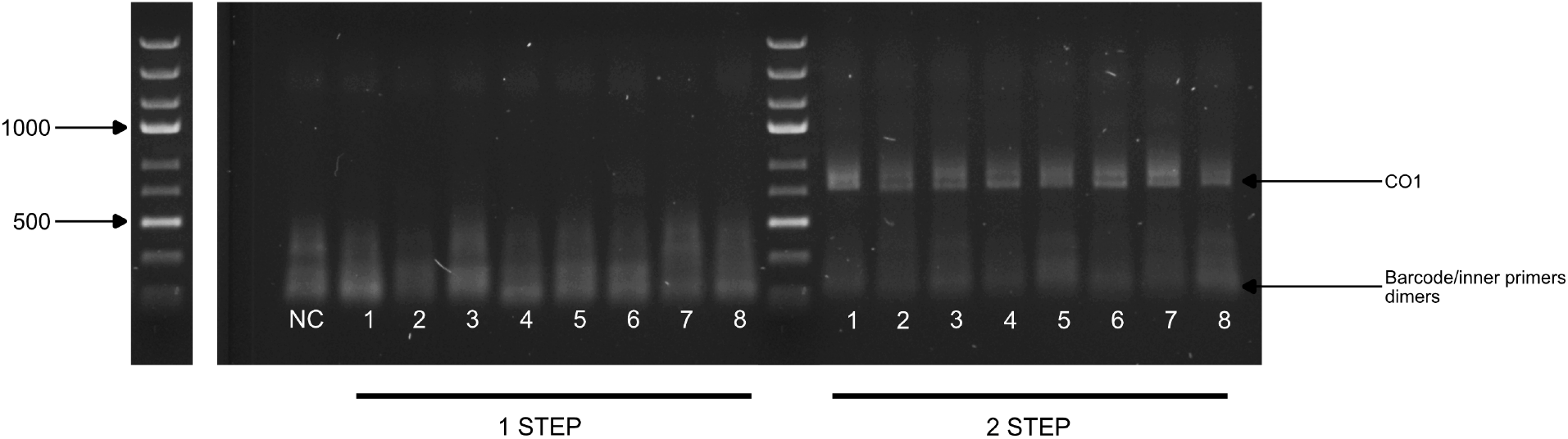
PCR comparison between the two-step and one-step protocols for the CO1 marker. Samples are numbered from 1 to 8, each representing a different specimen of Asellus aquaticus. Lane M: molecular size markers. Molecular size (in base pairs) is indicated on the left.

## Conclusion

We compared 4 primers PCR protocol initially developped by Matsuo et al., 2021 with and without the “drop in the lid” method developed by Macedo Rafael De Arruda et al., 2024. We amplified 2 different markers, CO1 and ITS on *Asellus*.*sp* DNA extract. The results showed a significant improvement of PCR yield when using the “drop in the lid” method displaying the possibility to perform barcoding PCR in a single PCR cycle wihtout the necessity to directly interact with the PCR’s master mix. This method allow to greatly reduce the risk of cross-contamination as well as reducing the time needed to perform DNA (meta)barcoding. Furthermore, using Nanopore sequencing for sequencing and Guppy for Basecalling, Demultiplexing, results show a 82% ratio of classified reads (results not displayed here), showing an efficient barcoding during the second step of the PCR. We hereby show that satisfactory PCR amplification can be achieved for a large variety of marker using a 4 primer PCR approach despite the unavoidable interaction between inner primers and barcodes, and that those results can be obtained without using deticated kit or expensive ligase.

## Data, script, code, and supplementary information availability

Sequences obtained through the experiment are available online at https://github.com/smartise/TWO-STEP_PCR

